# Optimal release of gene drives in population connectivity networks

**DOI:** 10.64898/2026.05.11.724203

**Authors:** Jonathan Halperin, Shachar Perlman, Shahar Shemesh, Keith D. Harris, Gili Greenbaum

**Author notes:** Contributed equally: Jonathan Halperin and Shachar Perlman.

## Abstract

Gene drives, genetic constructs that can spread deleterious alleles in wild populations, have the potential to address some of the major pressing challenges of the Anthropocene such as invasive species, spread of disease vectors, and agricultural pests. However, responsible and effective deployment of gene drive requires taking into account the complex nature of real-world population connectivity networks. In particular, it is unclear how the topological position of the deployment site affects the spread process and its final outcome. Here we develop a framework for modeling gene drive spread in population connectivity networks, and study the eco-evolutionary dynamics of gene drive spread under complex population structures. We investigated the relationship between the position of the deployment site in the topology of the network and whether the gene drive is eventually lost, fixed, or maintained at an intermediate frequency. We identified network centrality measures of deployment sites that are highly correlated with the outcome of deployment for different gene drive designs and across diverse network topologies. We also show that there is a trade-off between the time-to-fixation and the final outcome, implying that multiple centrality measures of the deployment site would need to be considered when aiming to achieve rapid and successful population control using gene drives.

## 1 Introduction

To address the growing ecological impacts of the Anthropocene, which amplify and globalize the burden of vector-born diseases [1, 2], agricultural pests [3], and invasive species [4, 5], novel approaches for population control are needed. One of the most promising technologies for suppression of harmful species is gene drives, where an artificial genetic construct is engineered to violate Mendelian inheritance, allowing it to rapidly spread and increase in frequency even under strong negative selection [6, 7, 8]. While lab experiments are useful for understanding basic behaviors of gene drives [9, 10, 11], because gene drives have yet to be tested in the field, studying their dynamics under natural ecological settings relies entirely on mathematical and computational models [12]. This is particularly true for the spatial dynamics of gene drive spread, which is a key feature that needs to be understood in order to informatively design and plan gene drive deployment schemes.

Whereas in a single panmictic population a gene drive is generally considered to have one of four different outcomes—fixation, loss, threshold-dependence (fixation above a frequency threshold and loss below it, i.e., unstable equilibrium), or persistence (stable equilibrium)— a wider range of outcomes need to be considered in spatially structured populations. For example, even when considering only two populations, gene drive deployment may result in differential targeting of one of the populations [13] or, when considering demography, oscillate in frequency or experience gene swamping [14, 15]. In continuous-space models, chasing dynamics may also emerge where the gene drive persists by allowing wild-type individuals to occupy empty areas [16, 17], and spatial Allee effects may lead to gene drive loss [12]. Some models aim to investigate the rate of spread of a successful drive [9, 17], whereas others are aimed at examining the ability of a gene drive to spread locally, avoiding spillover to non-target populations [13, 14]. Therefore, incorporating spatial spread in gene drive models reveals additional outcomes and adds considerations that need to be taken into account in gene drive development and deployment.

So far, gene drive models have incorporated simple and homogeneous spatial settings, such as two-deme models [13, 14], continuous space modeled using reaction-diffusion equations [18], and continuous space models using individual-based simulations [19]. However, natural population structures and connectivity patterns are typically heterogeneous, with different levels of connectivity between different locations. This connectivity heterogeneity may have critical effects on spread dynamics; for example, in epidemiology, considering the transmission connectivity patterns is crucial in order to understand whether a disease can spread in the population, and how this depends on the connectivity degree of an initial “patient zero” [20, 21, 22]. Therefore, in order to be applicable, gene drive models need to move beyond simplified spatial modeling and consider more complex connectivity patterns [12].

The most useful framework for studying spread dynamics under connectivity heterogeneity is network theory. In population networks, nodes represent populations, and edges represent migration connectivity (either at a fixed migration rate or using migration weights) [23, 24, 25]. In the context of population genetics, population networks have been used to study landscape connectivity and gene flow [26, 27], fragmentation [28], modular population structure [29, 30, 31], and the identification of dispersal pathways relevant to conservation and management [32, 33]. Using network theory is also important because it provides access to a wide toolkit of methodologies, such as generative models of random networks with different types of topological features [34], and centrality measures that capture different aspects of the topological position of a node in the network. Therefore, a population-network framework for studying gene drive spread may be useful for expanding the type of connectivity patterns that can be studied, and to bring gene drive models closer to the real-world systems where gene drives will potentially be deployed [12, 35].

When considering the non-homogeneous structure of population networks in the context of gene drives, important questions arise—how does the spatial position of the deployment site of the gene drive effect the spread dynamics and the final outcome? And, in cases when the outcome changes in relation to the deployment site, what information can be used to predict the outcome and the speed of gene drive spread? Answering these questions is crucial as gene drives approach testing in the wild, either in presumably locally contained field trials that are embedded in a larger metapopulation, or later in full releases in structured populations.

Here, we develop a flexible population-network framework for gene drive modeling under spatial heterogeneity, and use it to study the effect of the position of the release site on gene drive spread. We track spread dynamics in different networks and different release sites, and evaluate the association between the deployment site, its network properties, and the outcome and duration of the gene drive spread process. We focus on identifying network metrics that are correlated with key aspects of the process, such as eventual outcome and time of spread, so as to gain insight into what ecological connectivity features could be useful for evaluating risk and optimizing outcomes in gene drive projects.

## 2 Methods

To understand how population network structure influences the outcome of gene drive deployment, we developed a mathematical model of the dynamics of gene drive spread in population networks. We use this model in a simulation framework where we generate random population networks, randomly select a population in which the gene drive is deployed, and mathematically track the outcome of the spread of the gene drive. By analyzing the centrality of the population in which the gene drive is released, we study the effect of network topological properties on gene drive deployment outcomes and the time during which they occur.

### 2.1 Gene drive spread in population networks

To model gene drive dynamics in complex population structures, we extend a previously described two-population gene drive model [13] to apply to population networks. The model consists of two main steps: gene flow between populations, followed by selection and conversion within each population. We consider *n* populations, with migration occurring only between some pairs of connected populations. These gene flow dynamics can be described by a network with adjacency matrix *M* (henceforth ‘migration matrix’). We assume fixed and symmetric migration, i.e, *M*_*ij*_ = *M*_*ji*_ = *m* when populations *i* and *j* are connected, where *m* is in terms of the proportion of the population migrating.

To model the spread of a gene drive in a population network, we track the vector that describes the frequency of the gene drive in each of the populations through time, ***q***(*t*) = (*q*_1_(*t*), *q*_2_(*t*), …, *q*_*n*_(*t*))^⊤^. Initially, we consider that the gene drive is introduced to a single randomly chosen population *i* (the ‘*deployment population*’), so that *q*_*i*_(0) = 0.8 and *q*_*j*_(0) = 0 for all *j* ≠ *i*. The gene drive frequencies in each population after one generation of migration can be computed by taking the dot product of the migration matrix *M* with ***q***(*t*):

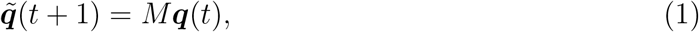

where 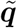 is the vector of gene drive frequencies after gene flow has occurred.

Next, we model the changes in the gene drive allele frequencies due to selection and the conversion of heterozygotes to gene drive homozygotes. We consider a standard gametic drive, which is characterized by the fitness of the gene drive allele *s*, the conversion probability *c*, and the dominance parameter of the gene drive allele *h* [6, 7]. For this, as in two-population models, we can use for each population standard equations that describe the effect of selection and conversion [13]:

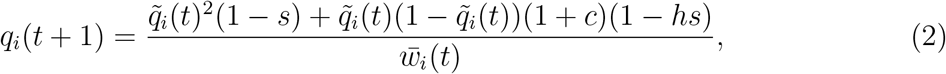

where 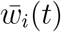 is the average fitness at time *t* and 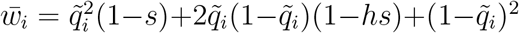. The two steps described in Eqs. 1 and 2 describe the change in frequencies from ***q***(*t*) to ***q***(*t* + 1); note that the order in which the two steps are implemented does not have a substantial impact on the resulting change in frequencies [13]. In all our analyses, we fixed the migration rate at *m* = 0.01 (See Supplemental Information for alternative parameterizations, with similar results).

### 2.2 Modeling population networks

To model different types of population structures, we generate random population networks using the random geometric graph (RGG) algorithm. This algorithm places *n* points uniformly at random in a Euclidean unit square, and connects all population pairs that are closer than a set distance parameter *d*. RGG population networks represent systems with distance-limited migration, which is typical in many real-world systems, where migration occurs primarily between geographically proximate populations [36, 23, 28]. In our simulations we used *n* = 100 and *d* = 0.175, which ensures network connectivity in RGG networks (two-dimensional RGG networks on a unit square are almost surely connected for 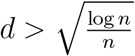 [37], with the threshold for *n* = 100 being 0.215); disconnected networks, in the cases that they were generated (about 15% of the simulations), were discarded and re-generated. For comparison and evaluation of the robustness of our results to different network structures, we also simulated networks with different structural properties using two commonly used generative network algorithms, Erdős–Rényi (ER) and Barabási–Albert (BA) [38, 39] (see Supplemental Information).

Because we incorporate heterogeneity in migration by assuming a network structure, there are different ways to consider the effect of a fixed migration parameter *m* on the migration network. Therefore, to understand how robust our results are to this modeling decision, we also evaluated alternative migration patterns where we fix either the overall incoming (‘fixed-in’) or outgoing (‘fixed-out’) migration from each population. Unlike our basic symmetric-migration model, fixing migration rates this way may induce asymmetry in migration rates along edges, and we therefore consider the network to be a directed network in these cases. In these two alternative models, migration rates do not scale directly with the degree of the nodes as in the symmetric model (where more nodes mean more incoming/outgoing migration), but rather assume that there is a fixed amount of individuals that can migrate out (for fixed-out) or be accepted into (for fixed-in) the population.

In each simulation, the deployment population was drawn randomly from the *n* populations, and we computed the centrality measures of the deployment population. We considered three well-studied centrality measures [40]: (i) *Degree centrality*, which measures the number of populations a focal population is connected to (i.e., the number of neighbors); (ii) *Eigenvector centrality*, which measures the influence of the focal population based both on its direct connectivity and the connectivity of its neighbors, capturing the role of highly connected hubs in shaping regional gene flow; (iii) *Closeness centrality*, which measures how efficiently a node can exchange migrants with all other nodes, calculated as the reciprocal of the sum of the length of all the geodesics between the node the other nodes in the network (i.e., populations with a high closeness centrality have, on average, the shortest distances to all other populations). Degree centrality captures local features of the network in the vicinity of the deployment population, while eigenvector and closeness centrality capture features relating to the position of the deployment population in the context of the entire topology of the network. While there exists a correlation between these measures in some types of networks, each measure captures a qualitatively different aspect of the topology of the population network. In each simulation, we computed the three centrality measures of the deployment population using the *igraph* Python package [41].

To evaluate the effect of the centrality of the deployment population on gene drive spread, we generated *N* = 250000 simulation repeats for each gene drive design (weak and strong drives), where each repeat included generation of a population network, random selection of the deployment population, and a deterministic computation of the spread of the gene drive. We simulated population networks using the RGG algorithm, or alternatively the ER or BA algorithms. The parameters of the algorithms were set so that the average degree of the generated networks would be similar: distance threshold *d* = 0.175 for RGG, per-dyad probability of connectivity *p* = 0.1 for ER, and the number of edges added during every iteration of graph generation *m* = 2 for BA.

### 2.3 Classifying gene drive spread outcomes

While in the two-population setting it is clear how to classify outcomes—fixation of the gene drive in the two populations, loss of the drive, stable polymorphism, and differential-targeting where the gene drive persists in high frequency in the deployment population and in low frequency in the other population [13]—classification of outcomes in a multi-population context is more challenging. We define three outcomes of interest: (i) ‘fixation’, defined as having the gene drive spread to frequency above 0.9 in 80% of the populations after 1000 generations; (ii) ‘failure’, defined as having the gene-drive allele frequency be below 0.1 in all populations after 1000 generations; (iii) ‘persistence’, in all other cases. This classification of outcomes allows us to distinguish between cases where the gene drive is expected to have a large global effect and likely suppress the entire population network (fixation), where it is likely to have little or no affect (failure), and where it is likely to affected only a limited part of the network (persistence). The persistence outcome may, in principle, also be observed in cases where the gene drive configuration is such that it generates a stable equilibrium [6, 7], but these equilibria do not exist within the range of gene drive parameters we used throughout our analyses [13]. Note that the simulation run-time of 1000 generations was selected to make sure that dynamics that approach fixation or failure are indeed classified as such, and therefore the ‘persistence’ outcome captures dynamics where there is a stable state in which the gene drive persists long-term. In practice, failure and fixation outcomes typically relate to simulations that rapidly reached a fixed state, either *q*_*i*_ = 0 or *q*_*i*_ = 1 in all nodes, respectively.

## 3 Results

We observe that the position of the deployment population in the population network can affect the dynamics of gene drive spread and the resulting outcomes. For example, considering a gene drive with a moderately high fitness cost (*s* = 0.55, *h* = 1, *c* = 1; ‘strong gene drive’), we observe that if the gene drive is deployed in a certain position, it spreads and rapidly fixes in the entire population network (fixation, Fig. 1A panels (ii) and (iii)). We also observe scenarios where the gene drive is deployed in another node and spreads, but does not take over the entire population (persistence, Fig. 1A panel (i)). Moreover, even when the gene drive does spread to reach global fixation, the rate at which this process occurs depends on the position of the deployment population (Fig. 1B). When considering a gene drive with a higher fitness cost (*s* = 0.65, *h* = 1, *c* = 1; ‘weak gene drive’) on the same network, we also observe that the outcome depends on the position of the deployment population: deployment in some positions results in failure and global loss of the gene drive, whereas deployment in other positions can result in persistence of the gene drive in a single population (Fig. 1C and D). These examples also illustrate that the deployment outcome still depends on the fitness cost of the gene drive as well, with deployment in the same population resulting in failure for the weak gene drive and global fixation for the strong gene drive (green arrow in Fig. 1A and C). Likewise, the difference in the fitness cost of the gene drive resulted in limited spread to a single population for the weak gene drive and extensive spread for the strong gene drive (purple and red arrows in Fig. 1A and C). Therefore, in order to understand the outcome and the dynamics of gene drive spread, the design parameters of the gene drive together with the topological position of the deployment population must be taken into account. For tractability, we compare these two parameterization of ‘weak’ and ‘strong’ gene drives in all analyses below.

**Figure 1:**
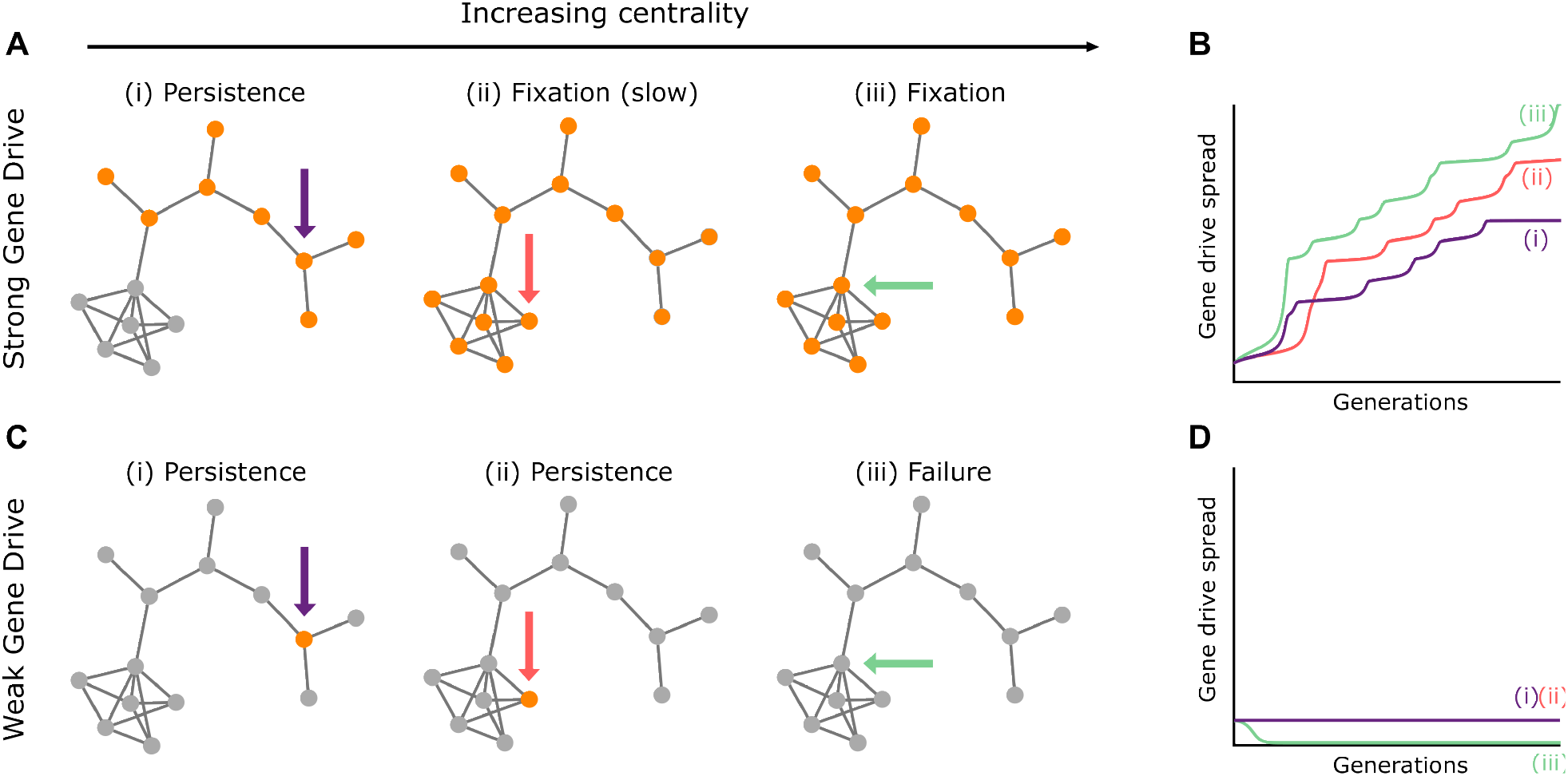
Relationship between the position of gene drive deployment position and outcome in a population network. Shown is an illustrative example on a single population network (nodes denote populations, edges denote migration connectivity). (A) A “strong” gene drive (*s* = 0.55, *c* = 1, *h* = 1) is deployed at three different positions with increasing centrality (black arrow at the top), indicated by colored arrows. Nodes colored in gray denote eventual loss or low frequency of the drive allele, and nodes in orange denote population where the gene drive eventually fixes or reaches very high frequency. The outcome category is denoted above each network. (B) The rate of spread of the gene drive over 100 generations with ‘strong” gene drive deployment, for the three deployment locations. (C) and (D) show the same analysis as panels (A) and (B), except that a “weak” gene drive is deployed (*s* = 0.65, *c* = 1, *h* = 1).

### 3.1 Centrality of the deployment population affects dynamics and outcomes

To understand how the topological position of the deployment population affects the outcome of deployment, we tracked its centrality using three centrality measures, each capturing different features of the topology of the population network. For each value of the centrality measure, we calculated the proportion of simulations that reached each outcome (Fig. 2). This analysis produces estimates of the probability that a deployment scenario initiated at a population with a given centrality value results in each of the outcomes. For example, deploying the strong gene drive in a deployment population with degree centrality of 2 is slightly more likely to result in persistence (54%) than fixation (46%) (Fig. 2A), whereas for the weak gene drive this would almost certainly result in persistence of the gene drive (Fig. 2D).

**Figure 2:**
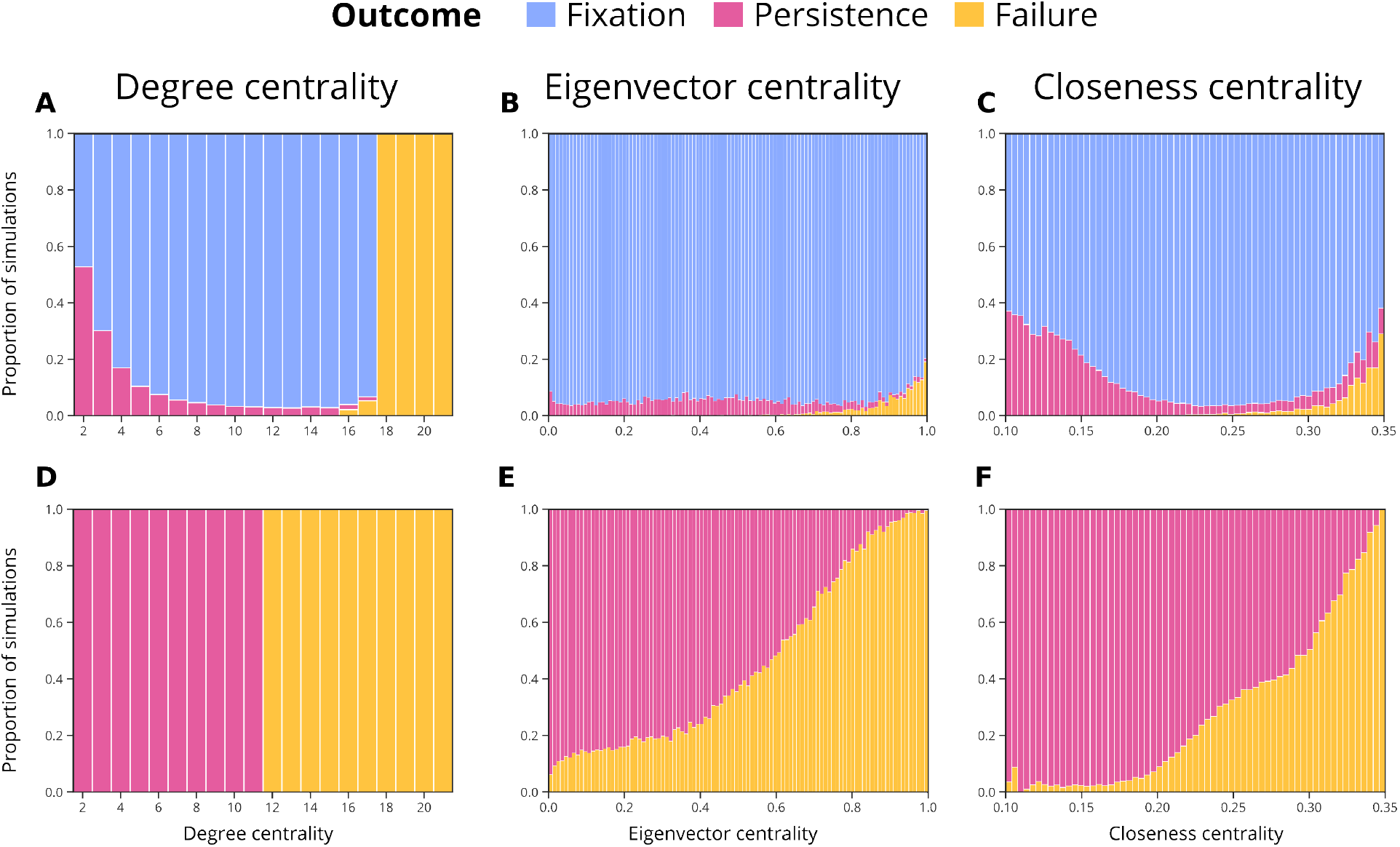
Relationship between centrality measures of the deployment population and deployment outcomes. Each panel shows results from *N* = 256000 simulations of random networks and deployment positions (RGG networks with parameters *n* = 100 and *r* = 0.175), run for *t* = 1000 generations. For each centrality value bin (numbers on the x-axis indicate the upper bound of each bin), the proportion of simulations reaching each of the outcomes is shown. (A)–(C) Results for a strong gene drive configuration with different centrality measures (degree, eigenvector and closeness centrality); (D)–(F) results for a weak gene drive.

When considering these three centrality measures, we observe that degree centrality is highly predictive of the outcome of deployment. For the strong gene drive, for low degree centrality scores the outcome is either persistence or fixation, with fixation becoming more likely at intermediate centrality scores (Fig. 2A). However, when the degree centrality scores cross a threshold (*k* = 18 for the strong drive and *k* = 12 for the weak drive; Fig. 2A and D), the outcome is almost certainly failure. This failure threshold emerges because releasing a gene drive into a population with high degree means that, initially, the incoming gene flow of wild-type alleles generates significant gene swamping that rapidly leads to the loss of the gene drive allele [14]. This degree-threshold emerges at the critical value for gene swamping to cause failure, as migration out of a population scales with its degree in this migration model (but not in our alternative migration models). Therefore, for this migration model, degree centrality is expected to be strongly correlated with the outcome of deployment. Note that the persistence observed with the weak gene drive is limited to the deployment population; this resembles the ‘differential targeting’ outcome described in a two-population model [13]. In this outcome, the gene drive spreads to high frequencies only in the target population, as opposed to persistence with the strong gene drive, which encompasses a wide range of levels of spread, i.e. a variable number of populations the gene drive has become fixed in.

In our robustness analyses, we observed qualitatively similar results when analyzing network types other than RGG (ER and BA networks); however, some outcomes were more or less frequent with respect to the centrality score compared to RGG (Fig. S1 and S2). For the strong drive scenario, persistence was more often observed in the ER network compared with RGG and BA networks, and eventual fixation was rare in the ER network (Figs. 2A–C, S1A–C and S2A–C). With the weak gene drive, the behavior was similar in all three models (Figs. 2D–F, S1D–F and S2D–F).

When applying our alternative migration models (fixed-in and fixed-out migration), we observe qualitatively similar results as with the original symmetric non-fixed model (Fig. S3 and S4). Importantly, migration in our alternative migration models does not linearly scale with population degree as in our main model, which indicates that our results are not a result of this scaling.

### 3.2 Association between deployment population centrality scores and outcomes

To understand which topological aspects of the population network are important for predicting that a certain outcome will occur, we analyzed the contribution of the deployment population centrality scores to the probability of occurrence of different outcomes using receiver-operating characteristic (ROC) curves. These ROC curves measure how well each centrality measure can be used to classify or predict the outcome, taking into account true- and false-positive rates across all decision thresholds [42]. For each curve, we computed the area-under-curve (AUC) value, which summarizes the accuracy of the centrality measure’s classification of the outcome. We analyzed the ROC curves of the measures with respect to two outcomes: (i) persistence with the strong gene drive, and (ii) failure with the weak gene drive. These two outcomes are considered to be undesired for different scenarios: persistence when the intention is for a strong drive to spread throughout the network, and failure when the intention is for a weak gene drive to spread in a target population.

In both cases, we observe that degree centrality classifies the outcome with the highest accuracy (Fig. 3). In the case of failure classification with the weak gene drive, degree centrality is a perfect classifier with AUC value equal to *AUC* = 1.0 (Fig. 3B), which reflects the degree-threshold generated by gene swamping that we observed in Figure 2D. Eigenvector centrality and closeness centrality are also correlated with failure, with the probability of failure increasing with higher values of both measures (Fig. 2B–C,E–F). This trend is also observed when using alternative network models, with a correlation between all three measures and the outcome of deployment (Fig. S5 and S6). As in the RGG model, we see that degree centrality is a perfect classifier of the failure outcome with the weak gene drive, whereas the correlations with the other two measures are weaker.

**Figure 3:**
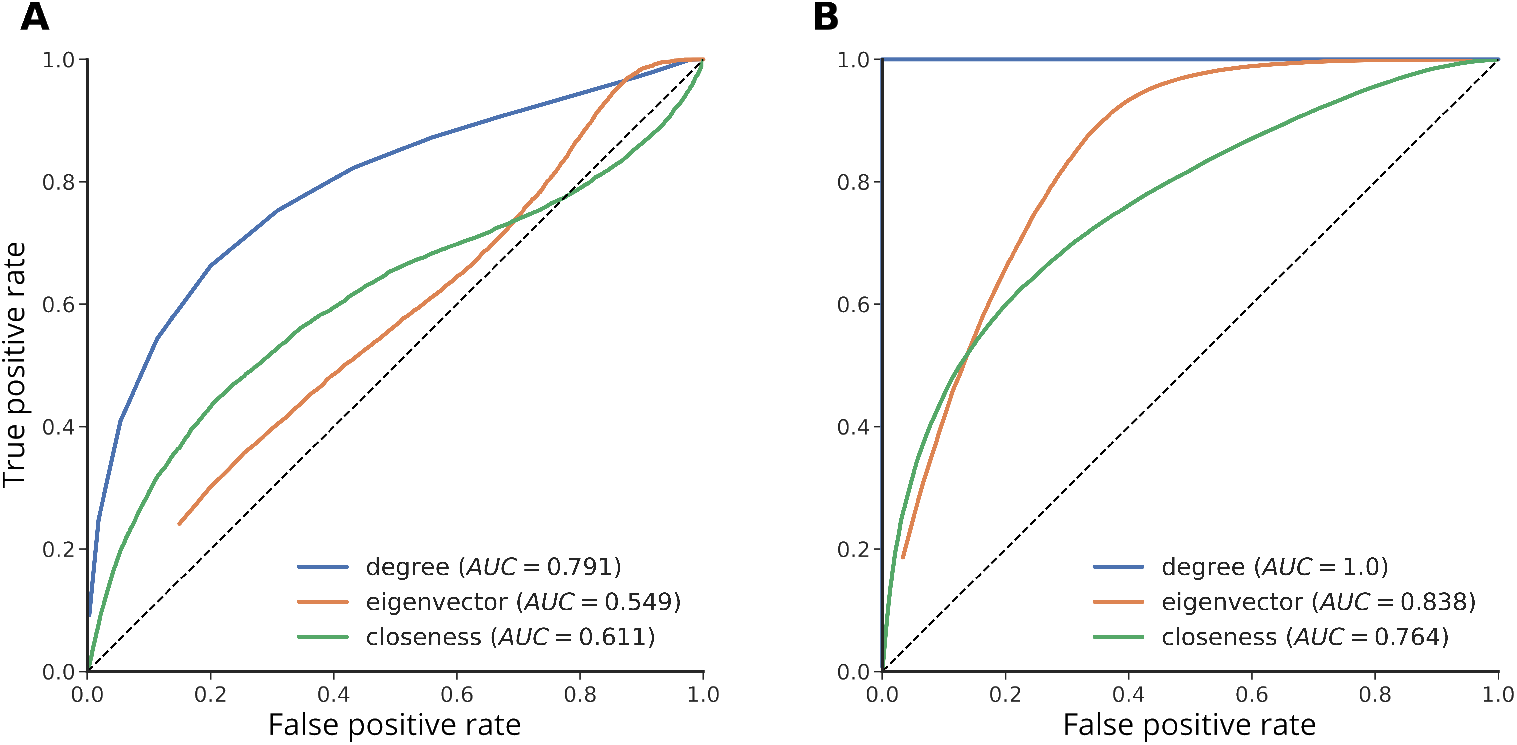
Classification and prediction of the outcomes based on the centrality of the deployment population. Shown are receiver-operating characteristic (ROC) curves, which quantify the contribution of each centrality measure to the probability that a certain outcome occurs. The area-under-curve (AUC) value quantifies the total contribution of a measure to a numerical value, where *AUC* = 0.5 denotes a random classifier (shown as the dotted line), and *AUC* = 1 denotes a perfect classifier. (A) The probability to correctly classify ‘persistence’ using the three centrality measures, with the ‘strong’ gene drive design. (B) The probability to correctly classify ‘failure’ using the three centrality measures, with the ‘weak’ gene drive design.

### 3.3 The time-to-fixation is shorter for closeness-central deployment populations

While achieving the desired outcome could be considered as the main measure of success of a deployment process, the number of generations to reach this outcome should also be considered. Minimizing this duration is important because, as the drive spreads through the population, unwanted consequences such as evolution of resistance to the gene drive and spillovers (to other species or population networks) may occur. Therefore, in order to reduce the risks involved, the time-to-fixation should be minimized. To investigate the association between the position of the deployment population and time-to-fixation, we studied a drive with an even lower fitness cost than our ‘strong drive’ (*s* = 0.4), so as to ensure that all outcomes attained are ‘fixation’.

We find that the centrality measures of the deployment population are indeed significantly associated with the time-to-fixation, with deployment in more central populations resulting in more rapid fixation (Fig. 4). Degree centrality of the deployment populationwas moderately correlated with time-to-fixation (*r*^2^ = 0.167), with a reduction of about 37% in time-to-fixation when comparing deployment in a population with degree 2 to one with degree 20 (Fig. 4A). This is likely due to the local ‘boost’ in the spread process that high degree populations gain at the early stage of the spread process. For the more global measure of closeness centrality, however, this association was much stronger (*r*^2^ = 0.798), and the reduction in the time to-fixation between the lower and higher centrality values was almost 70% (Fig. 4C). This is likely due to the shorter distances that the drive needs to spread when deployed in a closeness-central node to reach all peripheral populations, compared to a low closeness centrality deployment, which likely takes longer due to distant peripheral pockets that remain unaffected for longer. Similar results were observed with alternative migration models (Fig. S7).

**Figure 4:**
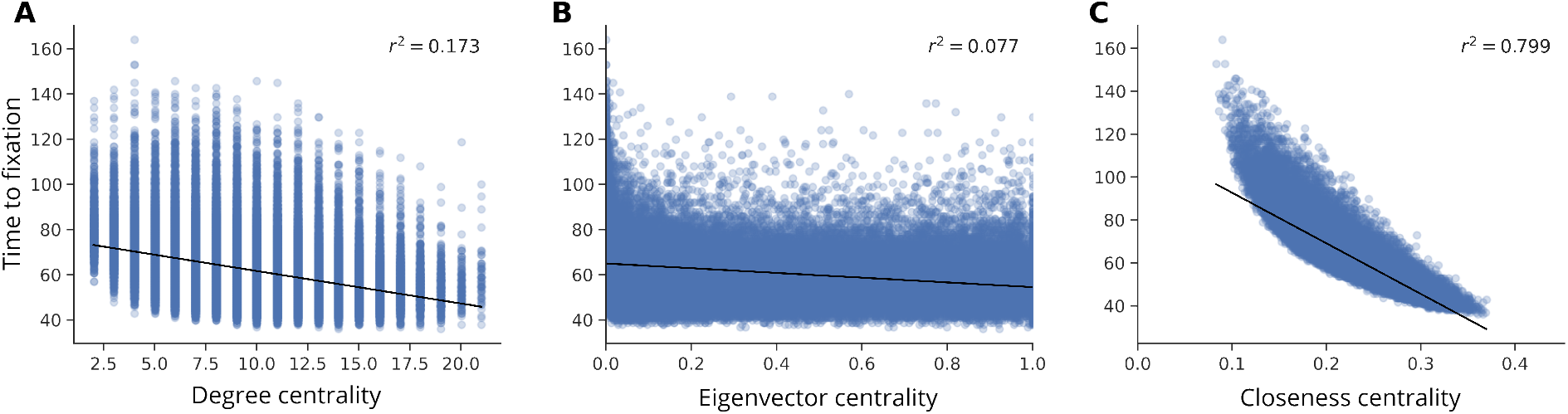
Time-to-fixation for gene drive deployments at populations with different centrality scores. The time-to-fixation is defined as the number of generations until the gene drive frequency in the entire population network is ≥ 0.9. Each dot is the time to fixation in a single simulation, and the x-axis shows the centrality value of the deployment population in that simulation. The black lines denote the linear regression of time-to-fixation values, and the coefficient of determination (*r*^2^) is shown at the top of each panel. (A) Degree centrality of the deployment population. (B) Eigenvector centrality of the deployment population. (C) Closeness centrality of the deployment population.

## 4 Discussion

We developed a mathematical framework for modeling gene drive spread in arbitrary networks by extending formulations of two-deme models. Using this framework, we showed that the position of the deployment population of the gene drive in the network can dramatically affect the outcome and the duration of the gene drive spread process. We also show that different centrality measures of the deployment population can be informative on the probability of attaining each of the outcomes and on the expected time-to-fixation. These results highlight the importance of considering the locations of gene drive releases in the context of the entire connectivity network when planning gene drive projects.

Our results suggest that obtaining ecological information, either locally (in the neighborhood of the considered deployment site) or globally (in the entire metapopulation) may be crucial for evaluating the success of the project. For example, when searching for a site for a field trial, conducting preliminary ecological studies on population connectivity of the site (i.e., its degree centrality) is crucial in order to maximize the probability of success. In this case, a weak drive is likely to be deployed, and we should prefer low-degree populations that are less likely to result in spillover or loss (Fig. 2A). If our goal is for the gene drive to spread in the entire network, however, we should prefer intermediate-degree deployment populations for the weak drive (Fig. 2A), and low-degree deployment populations with the strong drive (Fig. 2D), emphasizing that the design of the gene drive (e.g., its fitness cost) interacts with the position of the deployment population to generate the outcome.

Our finding that local connectivity is the main predictor for the outcome of the process (Fig. 3) is encouraging, because it suggests that it is not necessary to understand the entire connectivity network in order to coherently evaluate the likelihood of success, and that focused field surveys around the site of release may be sufficient. This was strongly evident when trying to predict the failure outcome, where degree was a perfect predictor (Fig. 3B), but less for predicting persistence. This is likely because determining persistence relies more heavily on connectivity patterns far from the deployment population (e.g., a narrow “bridge in the network” may halt the spread of the gene drive), whereas failure usually occurs due to gene swamping at the initial release location. However, because in many cases the expected duration of the process is also a key feature that will go into planning considerations, either due to risks of evolution of resistance [43] and spillovers [13, 44], or for developing a post-release monitoring program [45, 46], information on the entire connectivity network may be useful. We examined here only three centrality measures, but this limited investigation indicates that measuring the closeness centrality of the intended deployment site—how peripheral or central it is in the context of the entire network—provides substantial information on the expected time-to-fixation of the gene drive. This indicates that there may be tradeoffs that would need to be considered when selecting release sites, because optimizing both closeness and degree centrality of the deployment population would not always be possible. Other centrality measures may provide additional information on the time-to-fixation or other aspects of the gene drive spread process, potentially introducing additional trade-offs.

The extension of demic gene drive modeling framework to network models allowed us to address questions relating to optimal release and the ecological connectivity properties of the deployment site. This extension is important because it opens new avenues for asking and answering questions regarding the effect of connectivity heterogeneity on gene drive spread: Are different gene drive constructs affected differently by spatial connectivity configurations? Can ecological interventions of connectivity increase or decrease improve the likelihood of success? If release in several sites is possible, what release strategy would be optimal in different population networks (see [17])? In addition to enabling the qualitative theoretical study of these questions, expanding the scope of gene drive models to networks is also important for providing information on specific gene drive projects, where ecological and genetic data can be assembled to construct population networks for the target system [47, 48]. For example, once an explicit population network structure has been ascertained, simulators (e.g., MGDrivE [35, 49, 50]) can be used to explore the interaction of many ecological features with the connectivity patterns, providing detailed information for risk assessment, deployment strategy design and monitoring protocols.

By incorporating ecological realism more fully into gene drive models, we can better understand how gene drives are likely to behave in natural populations and generate predictions that are relevant for both developers and regulators. We showed that heterogeneous connectivity alone can substantially alter both the outcome and the timescale of spread, and that simple descriptors of release site connectivity and position in the metapopulation can provide useful guidance for deployment planning. Bridging the gap between theory and real-world implementation is becoming increasingly important as gene drive research moves closer to field testing, and population network models can help bridge this gap. Developing modeling frameworks that can explicitly account for release site choice, heterogeneous connectivity, and other ecologically grounded features will therefore be essential for informing risk assessment, trial design, and eventual deployment decisions.

## 5 Data availability

An interactive version of the model is available through the `modelRxiv` repository [51] at https://modelrxiv.org/model/kzCgnD.

## References

[1] N. Vora, “Impact of anthropogenic environmental alterations on vector-borne diseases,” The medscape journal of medicine, vol. 10, no. 10, p. 238, 2008.

[2] W. M. De Souza and S. C. Weaver, “Effects of climate change and human activities on vector-borne diseases,” Nature reviews microbiology, vol. 22, no. 8, pp. 476–491, 2024.

[3] S. Savary, L. Willocquet, S. J. Pethybridge, P. Esker, N. McRoberts, and A. Nelson, “The global burden of pathogens and pests on major food crops,” Nature ecology & evolution, vol. 3, no. 3, pp. 430–439, 2019.

[4] R. Early, B. A. Bradley, J. S. Dukes, J. J. Lawler, J. D. Olden, D. M. Blumenthal, P. Gonzalez, E. D. Grosholz, I. Ibañez, L. P. Miller, et al., “Global threats from invasive alien species in the twenty-first century and national response capacities,” Nature communications, vol. 7, no. 1, p. 12485, 2016.

[5] H. Seebens, T. Blackburn, E. Dyer, P. Genovesi, P. Hulme, J. Jeschke, S. Pagad, P. Pyšek, M. Winter, M. Arianoutsou, et al., “No saturation in the accumulation of alien species worldwide. nat commun 8: 14435,” 2017.

[6] A. Deredec, A. Burt, and H. C. J. Godfray, “The population genetics of using homing endonuclease genes in vector and pest management,” Genetics, vol. 179, no. 4, pp. 2013–2026, 2008.

[7] R. L. Unckless, P. W. Messer, T. Connallon, and A. G. Clark, “Modeling the manipulation of natural populations by the mutagenic chain reaction,” Genetics, vol. 201, no. 2, pp. 425–431, 2015.

[8] A. Burt and V. Koufopanou, “Homing endonuclease genes: the rise and fall and rise again of a selfish element,” Current opinion in genetics & development, vol. 14, no. 6, pp. 609–615, 2004.

[9] K. Kyrou, A. M. Hammond, R. Galizi, N. Kranjc, A. Burt, A. K. Beaghton, T. Nolan, and A. Crisanti, “A crispr–cas9 gene drive targeting doublesex causes complete population suppression in caged *Anopheles gambiae* mosquitoes,” Nature Biotechnology, vol. 36, no. 11, pp. 1062–1066, 2018.

[10] J. Champer, E. Yang, E. Lee, J. Liu, A. G. Clark, and P. W. Messer, “A crispr homing gene drive targeting a haplolethal gene removes resistance alleles and successfully spreads through a cage population,” Proceedings of the National Academy of Sciences, vol. 117, no. 39, pp. 24377–24383, 2020.

[11] V. M. Gantz, N. Jasinskiene, O. Tatarenkova, A. Fazekas, V. M. Macias, E. Bier, and A. James, “Highly efficient cas9-mediated gene drive for population modification of the malaria vector mosquito *Anopheles stephensi*,” Proceedings of the National Academy of Sciences, vol. 112, no. 49, pp. E6736–E6743, 2015.

[12] J. Kim, K. D. Harris, I. K. Kim, S. Shemesh, P. W. Messer, and G. Greenbaum, “Incorporating ecology into gene drive modelling,” Ecology Letters, vol. 26, pp. S62–S80, 2023.

[13] G. Greenbaum, M. W. Feldman, N. A. Rosenberg, and J. Kim, “Designing gene drives to limit spillover to non-target populations,” PLoS Genetics, vol. 17, no. 2, p. e1009278, 2021.

[14] K. D. Harris and G. Greenbaum, “Rescue by gene swamping as a gene drive deployment strategy,” Cell Reports, vol. 42, no. 12, 2023.

[15] Z. Wen, M. Wan, G. Greenbaum, and O. Carja, “Mapping gene drive dynamics onto mendelian models,” bioRxiv, pp. 2026–01, 2026.

[16] J. Champer, I. K. Kim, S. E. Champer, A. G. Clark, and P. W. Messer, “Suppression gene drive in continuous space can result in unstable persistence of both drive and wild-type alleles,” Molecular Ecology, vol. 30, no. 4, pp. 1086–1101, 2021.

[17] Z. Wang and J. Champer, “Optimal spatial release strategies for confined gene drives and wolbachia,” bioRxiv, pp. 2026–03, 2026.

[18] L. Girardin and F. Débarre, “Demographic feedbacks can hamper the spatial spread of a gene drive,” Journal of Mathematical Biology, vol. 83, no. 6, pp. 1–33, 2021.

[19] J. Champer, I. K. Kim, S. E. Champer, A. G. Clark, and P. W. Messer, “Suppression gene drive in continuous space can result in unstable persistence of both drive and wild-type alleles,” Molecular Ecology, vol. 30, pp. 1086––1101, 2021.

[20] M. Kitsak, L. K. Gallos, S. Havlin, F. Liljeros, L. Muchnik, H. E. Stanley, and H. A. Makse, “Identification of influential spreaders in complex networks,” Nature physics, vol. 6, no. 11, pp. 888–893, 2010.

[21] L. Danon, A. P. Ford, T. House, C. P. Jewell, M. J. Keeling, G. O. Roberts, J. V. Ross, and M. C. Vernon, “Networks and the epidemiology of infectious disease,” Interdisciplinary perspectives on infectious diseases, vol. 2011, no. 1, p. 284909, 2011.

[22] D. Bucur and P. Holme, “Beyond ranking nodes: Predicting epidemic outbreak sizes by network centralities,” PLoS computational biology, vol. 16, no. 7, p. e1008052, 2020.

[23] G. Greenbaum and N. H. Fefferman, “Application of network methods for understanding evolutionary dynamics in discrete habitats,” Molecular ecology, vol. 26, no. 11, pp. 2850–2863, 2017.

[24] A. F. Rozenfeld, S. Arnaud-Haond, E. Hernández-García, V. M. Eguíluz, E. A. Serrão, and C. M. Duarte, “Network analysis identifies weak and strong links in a metapopulation system,” Proceedings of the National Academy of Sciences, vol. 105, no. 48, pp. 18824–18829, 2008.

[25] R. J. Dyer and J. D. Nason, “Population graphs: the graph theoretic shape of genetic structure,” Molecular ecology, vol. 13, no. 7, pp. 1713–1727, 2004.

[26] C. J. Garroway, J. Bowman, D. Carr, and P. J. Wilson, “Applications of graph theory to landscape genetics,” Evolutionary Applications, vol. 1, no. 4, pp. 620–630, 2008.

[27] R. J. Dyer, J. D. Nason, and R. C. Garrick, “Landscape modelling of gene flow: improved power using conditional genetic distance derived from the topology of population networks,” Molecular ecology, vol. 19, no. 17, pp. 3746–3759, 2010.

[28] O. Peled, J. Kim, and G. Greenbaum, “Network-based genetic monitoring of landscape fragmentation,” Proceedings of the National Academy of Sciences, vol. 123, no. 8, p. e2515033123, 2026.

[29] W. E. Peterman, B. H. Ousterhout, T. L. Anderson, D. L. Drake, R. D. Semlitsch, and L. S. Eggert, “Assessing modularity in genetic networks to manage spatially structured metapopulations,” Ecosphere, vol. 7, no. 2, p. e01231, 2016.

[30] B. E. Reichert, R. J. Fletcher Jr, C. E. Cattau, and W. M. Kitchens, “Consistent scaling of population structure across landscapes despite intraspecific variation in movement and connectivity,” Journal of Animal Ecology, vol. 85, no. 6, pp. 1563–1573, 2016.

[31] Y. S. Rodger, G. Greenbaum, M. Silver, S. Bar-David, and G. Winters, “Detecting hierarchical levels of connectivity in a population of acacia tortilis at the northern edge of the species’ global distribution: Combining classical population genetics and network analyses,” PLoS One, vol. 13, no. 4, p. e0194901, 2018.

[32] M. S. O’Donnell, D. R. Edmunds, C. L. Aldridge, J. A. Heinrichs, A. P. Monroe, P. S. Coates, B. G. Prochazka, S. E. Hanser, and L. A. Wiechman, “Defining fine-scaled population structure among continuously distributed populations,” Methods in Ecology and Evolution, vol. 13, no. 10, pp. 2222–2235, 2022.

[33] E. Boulanger, A. Dalongeville, M. Andrello, D. Mouillot, and S. Manel, “Spatial graphs highlight how multi-generational dispersal shapes landscape genetic patterns,” Ecography, vol. 43, no. 8, pp. 1167–1179, 2020.

[34] E. Delmas, M. Besson, M.-H. Brice, L. A. Burkle, G. V. Dalla Riva, M.-J. Fortin, D. Gravel, P. R. Guimarães Jr, D. H. Hembry, E. A. Newman, et al., “Analysing ecological networks of species interactions,” Biological Reviews, vol. 94, no. 1, pp. 16–36, 2019.

[35] H. M. Sánchez C, S. L. Wu, J. B. Bennett, and J. M. Marshall, “Mgdrive: A modular simulation framework for the spread of gene drives through spatially explicit mosquito populations,” Methods in Ecology and Evolution, vol. 11, no. 2, pp. 229–239, 2020.

[36] P. A. Hancock, A. North, A. W. Leach, P. Winskill, A. C. Ghani, H. C. J. Godfray Burt, and J. D. Mumford, “The potential of gene drives in malaria vector species to control malaria in african environments,” Nature Communications, vol. 15, no. 1, p. 8976, 2024.

[37] M. Penrose, Random geometric graphs, vol. 5. OUP Oxford, 2003.

[38] P. Erdős, A. Rényi, et al., “On the evolution of random graphs,” Publ. Math. Inst. Hung. Acad. Sci, vol. 5, no. 1, pp. 17–60, 1960.

[39] A.-L. Barabási and R. Albert, “Emergence of scaling in random networks,” science, vol. 286, no. 5439, pp. 509–512, 1999.

[40] J. Golbeck, Analyzing the social web. Newnes, 2013.

[41] G. Csardi and T. Nepusz, “The igraph software package for complex network research,” InterJournal, vol. Complex Systems, p. 1695, 2006.

[42] J. A. Hanley and B. J. McNeil, “The meaning and use of the area under a receiver operating characteristic (roc) curve.,” Radiology, vol. 143, no. 1, pp. 29–36, 1982.

[43] R. L. Unckless, A. G. Clark, and P. W. Messer, “Evolution of resistance against crispr/cas9 gene drive,” Genetics, vol. 205, no. 2, pp. 827–841, 2017.

[44] S. R. Rybnikov, A. Lampert, and G. Greenbaum, “Integrating gene drives with established pest controls to mitigate spillover risk,” bioRxiv, pp. 2025–09, 2025.

[45] G. Rašić, N. F. Lobo, E. H. Jeffrey Gutiérrez, H. M. Sánchez C, and J. M. Marshall, “Monitoring needs for gene drive mosquito projects: lessons from vector control field trials and invasive species,” Frontiers in genetics, vol. 12, p. 780327, 2022.

[46] K. C. Long, L. Alphey, G. J. Annas, C. S. Bloss, K. J. Campbell, J. Champer, C.-H. Chen, A. Choudhary, G. M. Church, J. P. Collins, et al., “Core commitments for field trials of gene drive organisms,” Science, vol. 370, no. 6523, pp. 1417–1419, 2020.

[47] J. M. Marshall, S. Yang, J. B. Bennett, I. Filipović, and G. Rašić, “Spatial close-kin mark-recapture methods to estimate dispersal parameters and barrier strength for mosquitoes,” PLOS Computational Biology, vol. 21, no. 11, p. e1013713, 2025.

[48] G. Rašić and J. M. Marshall, “Integrating mosquito genomics into simulation modeling: opportunities for better-informed biocontrol,” Current opinion in insect science, vol. 73, p. 101456, 2026.

[49] S. L. Wu, J. B. Bennett, H. M. Sánchez C, A. J. Dolgert, T. M. León, and J. M. Marshall, “Mgdrive 2: A simulation framework for gene drive systems incorporating seasonality and epidemiological dynamics,” PLoS computational biology, vol. 17, no. 5, p. e1009030, 2021.

[50] A. Mondal, H. M. Sánchez C, and J. M. Marshall, “Mgdrive 3: A decoupled vector-human framework for epidemiological simulation of mosquito genetic control tools and their surveillance,” PLoS computational biology, vol. 20, no. 5, p. e1012133, 2024.

[51] K. D. Harris, G. Hadari, and G. Greenbaum, “modelRxiv: A platform for the dissemination and interactive display of models,” Ecology Letters, vol. 28, no. 1, p. e70042, 2025.

