## Supplemental Information for "Optimal release of gene drives in population connectivity networks"

### **Supplementary Information** — Optimal release of gene drives in population connectivity networks

#### Alternative network models

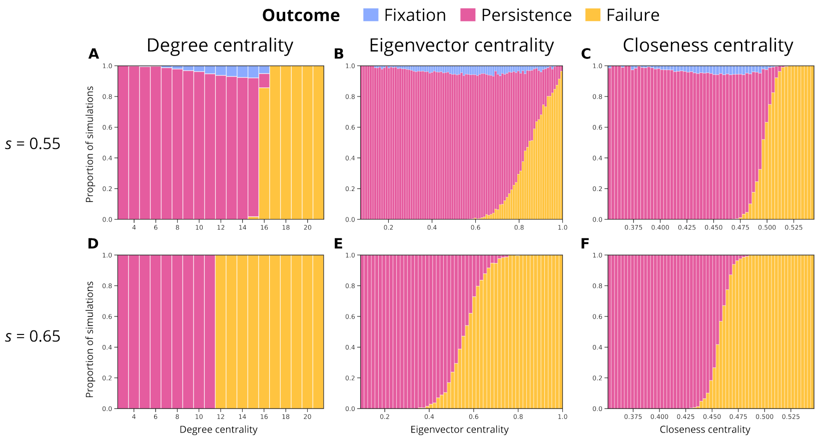


Figure. S1: **Relationship between centrality measures of the deployment population and outcomes for the Erdős–Rényi model.** This figure is the same as Figure 2 in the main text, except that the Erdős–Rényi model with parameters $n = 100$ and $p = 0.1$ was used instead of the RGG model. For each centrality value bin (numbers on the x-axis indicate the upper bound of each bin), the proportion of simulations reaching each of the outcomes is shown. (A)–(C) Results for a strong gene drive configuration with different centrality measures (degree, eigenvector and closeness centrality); (D)–(F) results for a weak gene drive.

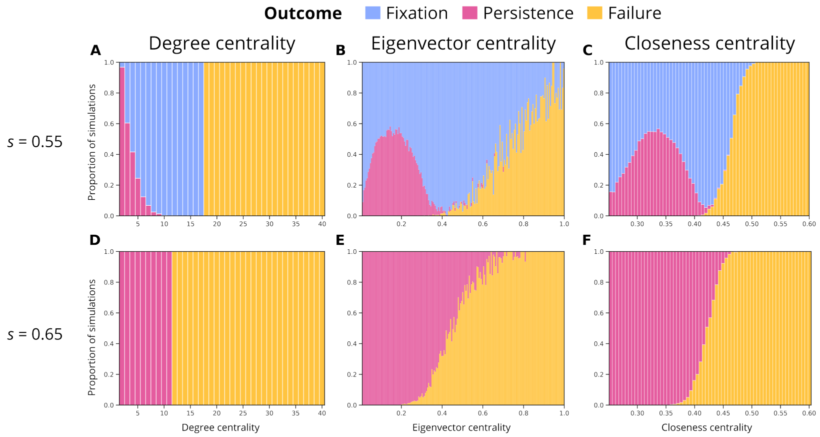


Figure. S2: **Relationship between centrality measures of the deployment population and outcomes for the Barabási–Albert model.** This figure is the same as Figure 2 in the main text, except that the Barabási–Albert model with parameters $n = 100$ and $m = 2$ was used instead of the RGG model. For each centrality value bin (numbers on the x-axis indicate the upper bound of each bin), the proportion of simulations reaching each of the outcomes is shown. (A)–(C) Results for a strong gene drive configuration with different centrality measures (degree, eigenvector and closeness centrality); (D)–(F) results for a weak gene drive.

### Alternative migration models

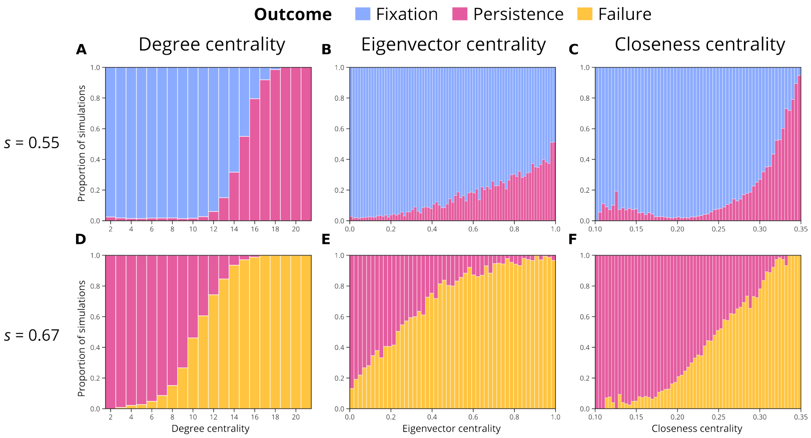


Figure. S3: **Relationship between centrality measures of the deployment population and outcomes for the fixed-in migration model.** This figure is the same as Figure 2 in the main text, except that the fixed-in migration mode (fixed overall incoming migration $m = 0.01$ for each population) was used instead of the RGG model. For each centrality value bin (numbers on the x-axis indicate the upper bound of each bin), the proportion of simulations reaching each of the outcomes is shown. (A)–(C) Results for a strong gene drive configuration with different centrality measures (degree, eigenvector and closeness centrality); (D)–(F) results for a weak gene drive.

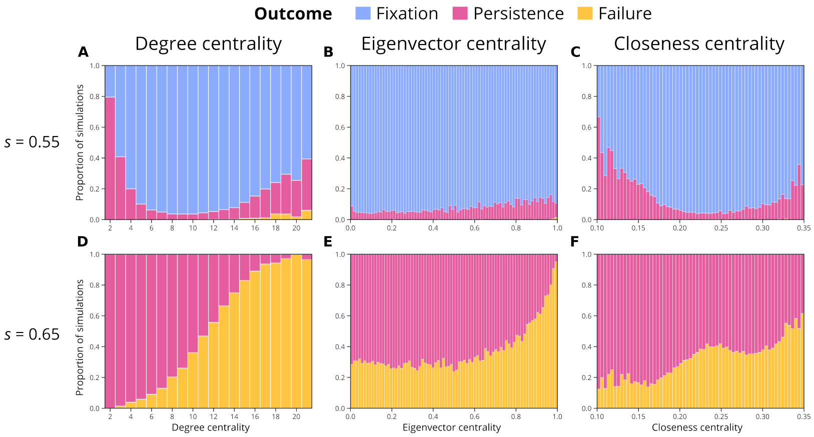


Figure. S4: **Relationship between centrality measures of the deployment population and outcomes for the fixed-out migration model.** This figure is the same as Figure 2 in the main text, except that the fixed-out migration mode (fixed overall outgoing migration $m = 0.01$ for each population) was used instead of the RGG model. For each centrality value bin (numbers on the x-axis indicate the upper bound of each bin), the proportion of simulations reaching each of the outcomes is shown. (A)–(C) Results for a strong gene drive configuration with different centrality measures (degree, eigenvector and closeness centrality); (D)–(F) results for a weak gene drive.

### Outcome prediction with alternative network models

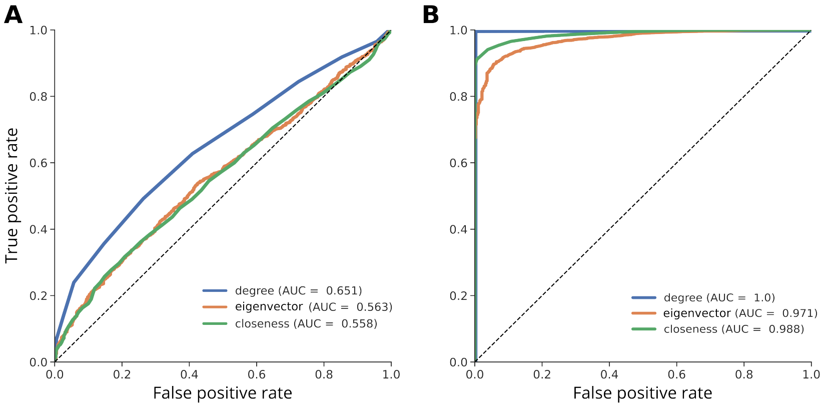


Figure S5: **Classification and prediction of the outcomes based on the centrality of the deployment population for the Erdős–Rényi model.** Shown are receiver-operating characteristic (ROC) curves and area-under-curve (AUC) values. Dotted line denoted the $AUC = 0.5$ random classifier. (A) The probability to correctly classify 'persistence' using the three centrality measures, with the 'strong' gene drive design. (B) The probability to correctly classify 'failure' using the three centrality measures, with the 'weak' gene drive design.

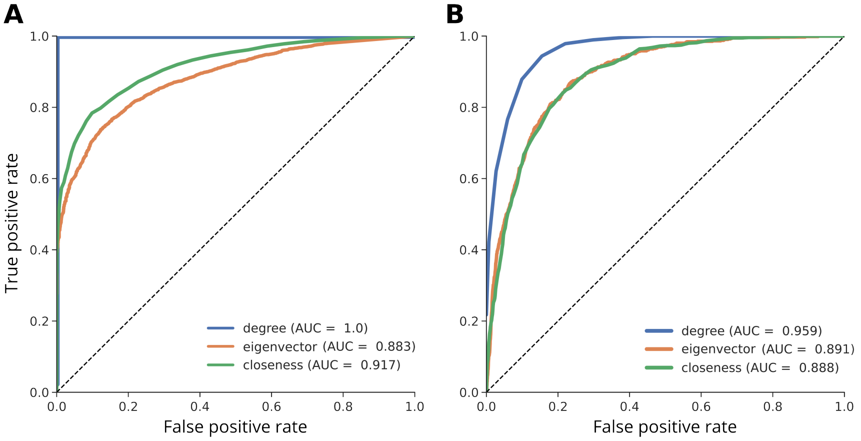


Figure S6: **Classification and prediction of the outcomes based on the centrality of the deployment population for the Barabási–Albert model.** Shown are receiver-operating characteristic (ROC) curves and area-under-curve (AUC) values. Dotted line denoted the $AUC = 0.5$ random classifier. (A) The probability to correctly classify 'persistence' using the three centrality measures, with the 'strong' gene drive design. (B) The probability to correctly classify 'failure' using the three centrality measures, with the 'weak' gene drive design.

### Time-to-fixation with alternative migration models

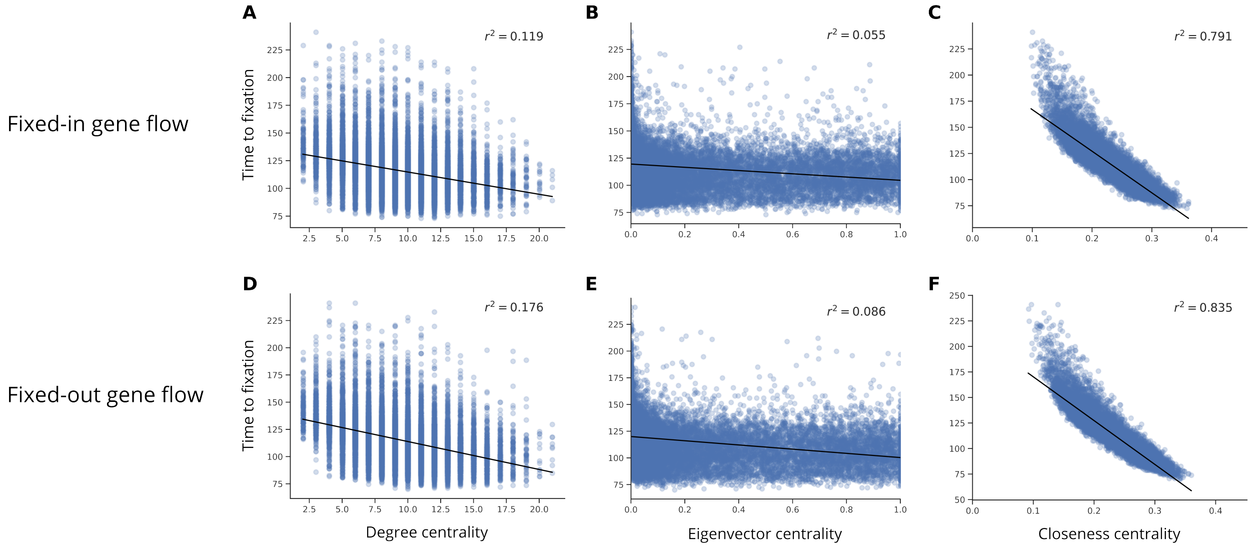


Figure. S7: **Time-to-fixation for gene drive deployments at populations with different centrality scores for alternative migration models.** The time-to-fixation is defined as the number of generations until the gene drive frequency in the entire population network is $\geq 0.9$. Each dot is the time to fixation in a single simulation, and the x-axis shows the centrality value of the deployment population in that simulation. The black lines denotes the linear regression of time-to-fixation values, and the coefficient of determination ($r^2$) is shown at the top of each panel. (A–C) Results for the fixed-in migration model. (D–F) Results for the fixed-in migration model.
